# Multiplexed dynamic imaging of genomic loci in single cells by combined CRISPR imaging and DNA sequential FISH

**DOI:** 10.1101/101477

**Authors:** Yodai Takei, Sheel Shah, Sho Harvey, Lei S. Qi, Long Cai

**Affiliations:** Division of Biology and Biological Engineering, California Institute of Technology, Pasadena, CA 91125, USA; UCLA-Caltech Medical Scientist Training Program, David Geffen School of Medicine, University of California at Los Angeles, Los Angeles, CA 90095, USA; Division of Chemistry and Chemical Engineering, California Institute of Technology, Pasadena, CA 91125, USA; Department of Bioengineering, Stanford University, Stanford, CA 94305, USA; Department of Chemical and Systems Biology, Stanford University, Stanford, CA 94305, USA; ChEM-H, Stanford University, Stanford, CA 94305, USA

## Abstract

Visualization of chromosome dynamics allows the investigation of spatiotemporal chromatin organization and its role in gene regulation and other cellular processes. However, current approaches to label multiple genomic loci in live cells have a fundamental limitation in the number of loci that can be labelled and uniquely identified. Here we describe an approach we call “track first and identify later” for multiplexed visualization of chromosome dynamics by combining two techniques: CRISPR labeling and DNA sequential fluorescence *in situ* hybridization (DNA seqFISH). Our approach first labels and tracks chromosomal loci in live cells with the CRISPR system, then barcodes those loci by DNA seqFISH in fixed cells and resolves their identities. We demonstrate our approach by tracking telomere dynamics, identifying 12 unique subtelomeric regions with variable detection efficiencies, and tracking back the telomere dynamics of respective chromosomes in mouse embryonic stem cells.

---

The three-dimensional chromatin organization in the nucleus plays an important role in gene regulation and other cellular processes (1,2). Visualizing spatiotemporal chromatin organization helps to interrogate its relationship with biological functions. Recently developed CRISPR systems can be a powerful and versatile tool to label and track genomic loci in live mammalian cells (3,4), supplementing dynamics to the static information from fluorescence *in situ* hybridization (FISH) in fixed cells. One of the challenges of live cell imaging of genomic loci is imaging multiple loci simultaneously in individual cells. To overcome this issue and enable multicolor CRISPR imaging, several methods have been developed by using orthogonal Cas9 systems (5,6) or engineered single guide RNA (sgRNA) scaffolds (7-9). However, even these methods only allow the simultaneous imaging of two or three loci. More recently, the color barcoding approach, using engineered sgRNA scaffolds recruiting different combinations of spectrally distinct fluorescent proteins, has demonstrated simultaneous imaging of six chromosomal loci in single cells (10). Although these multicolor approaches have expanded the potential of CRISPR imaging, they have a fundamental bottleneck in multiplexing due to the limited number of available orthogonal Cas9 systems, sgRNA scaffolds, or fluorescent proteins with spectrally distinct fluorophores.

Here we propose a new approach to label and distinguish multiple genomic loci using the combination of CRISPR imaging and DNA sequential FISH (DNA seqFISH), which provides large multiplexing capabilities. The principle of our approach is illustrated in Fig. 1. Multiple genomic loci are labeled with the CRISPR system all in a single color, and tracked in individual live cells. At the end of the live recording, cells are fixed and the identity of each locus is resolved by the color barcodes from DNA seqFISH. In this manner, even if the identities of labeled loci are indistinguishable during the live recording, as long as their positions are distinctly tracked in live imaging, these chromosomal loci can be subsequently identified with DNA seqFISH.

**FIGURE 1.**
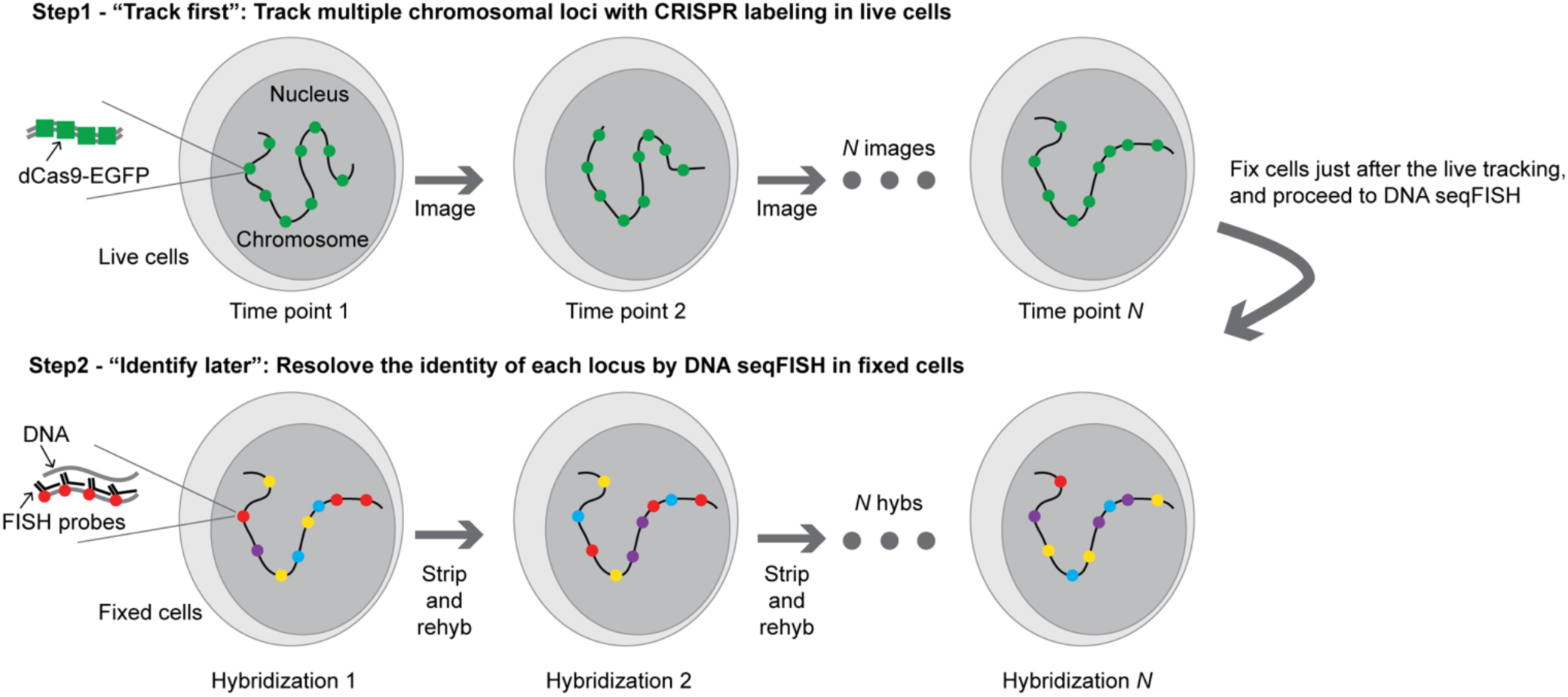
Schematic of the “track first and identify later” approach with the combination of the CRISPR labeling and DNA seqFISH techniques. Nine regions in one chromosome are illustrated in this schematic. Each chromosomal position can be identified from the DNA seqFISH step and its motion can be backtracked from the live imaging.

This “track first and identify later” approach can circumvent the multiplexing limitations of live cell imaging. As a proof-of-principle, we applied our technique to track telomeric loci in live mouse embryonic stem (mES) cells, and uniquely assigned 12 telomeric loci to particular chromosomes by performing DNA seqFISH of distal subtelomeric regions after the live tracking (Fig. 2 *A*).

**FIGURE 2.**
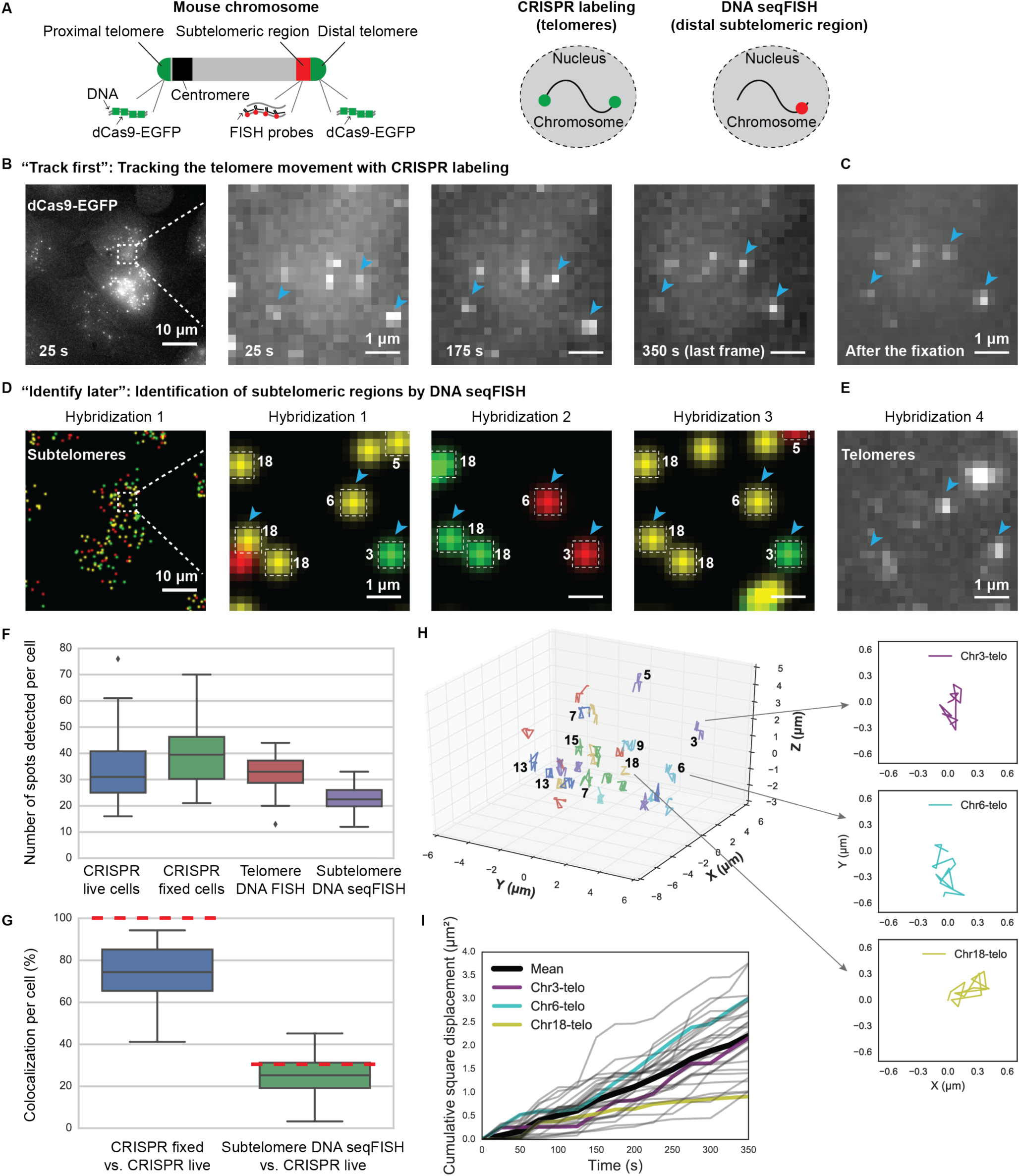
Multiplexed telomere tracking and identification of chromosomes with the "track first and identify later” approach in mES cells. (*A*) Schematic of the approach applied to telomere in a mouse chromosome. Proximal and distal telomere were labeled by the CRISPR system whereas only the distal subtelomeric region was labeled by DNA seqFISH. In total, 12 distal subtelomeric regions in 12 chromosomes were robustly read out by DNA seqFISH. (*B* and *C*) One-color telomere imaging in live cells at different time points (*B*) and after fixing cells (*C*), using the constructed mES cell line. (*D* and *E*) Composite digitized three-color (Alexa 647: red, Alexa 594: green and Cy3B: yellow) DNA seqFISH data for three rounds of hybridizations targeting subtelomeric regions (*D*), and one-color (Cy7) data for the fourth hybridization targeting telomeres (*E*). Based on the barcode identities, chromosome numbers are assigned to each of the subtelomeric spots (*D*). Note that DNA seqFISH spots do not perfectly colocalize with CRISPR imaging spots because they target adjacent regions in the genome. Dots without colocalization between hybridizations are due to nonspecific binding of probes or mis-hybridization in the cells. Images are maximum intensity projections of a z-stack of fluorescence images and the boxed region of the cell is magnified (*B-E*). (*F*) Comparing number of telomeric or subtelomeric spots detected per cell with the CRISPR and DNA seqFISH methods. In total, 938 CRISPR spots in live cells (last frame of the movie), 1138 CRISPR spots in fixed cells, 909 telomeric spots by DNA FISH and 628 subtelomeric spots by DNA seqFISH in 28 cells were analyzed. (*G*) Comparing colocalization percentage of spots detected per cell. Red dashed lines represent expected colocalization percentage per cell. (*H*) Trajectories of telomeric loci in the magnified cell. In this cell, 30 telomeric trajectories were detected from CRISPR imaging and 10 of these trajectories were uniquely assigned to particular chromosomes based on the subtelomere color barcodes. Trajectories of three loci in the magnified images (*B-E*) were also highlighted as xy projections (inset). Projected trajectories start from (0.0, 0.0). (*I*) Cumulative square displacement traces (*n* = 30) calculated with two adjacent frames as a function of time from the magnified cell. Traces of three loci in the magnified images (*B-E*) were shown as colored traces.

To observe the dynamics of telomeric loci in live mES cells, we generated a mES cell line stably expressing *Streptococcus pyogenes* nuclease-deactivated Cas9 (dCas9) fused to EGFP and sgRNA targeting telomeric loci by following a previous study (3). The dCas9 protein carried two nuclear localization signals for proper nuclear import. Using this clonal line, we performed live imaging over 6 min (Fig. 2 *B*, S1, Movie S1), and tracked the dynamics of telomeric loci in three-dimensional space.

Immediately after the live tracking, cells were fixed and processed for DNA seqFISH (Fig. 2 *B-E*). We quantified the number of telomeric dots (Fig. 2 *F*) and observed that on average, 73.0% of telomeric dots at the last frame of the live tracking were uniquely assigned to telomeric dots after the fixation (Fig. 2 *G*), indicating that the majority of the CRISPR labeled loci do not move significantly before and during fixing. Subtelomeric regions in respective chromosomes were barcoded based on a sequential barcoding method we demonstrated previously with RNA FISH (11,12). With this method, the number of loci that can be distinguished scales as *F^N^*, where *F* is the number of distinct fluorophores and *N* is the number of hybridization rounds. Each subtelomeric region was targeted with a set of FISH probes labeled with a single fluorophore during each round of hybridization. Specifically, the primary probes targeting the genomic loci also contain overhang sequences that are unique to each locus. A set of adapter probes that are dye labeled are hybridized to the overhang sequences (Fig. S2 *A*). We imaged cells, and then treated them with 70% formamide solution to displace the adapter probes (Fig. S2). We imaged cells again to confirm the probe displacement, and subsequent rounds of hybridizations were performed (Fig. S2 *B,C*). To cover 12 subtelomeric regions (Table S1), we used three dyes and three rounds of hybridizations (Fig. 2 *D*). We also used a fourth round of hybridization to image telomeres with DNA FISH (Fig. 2 *E*), and three different subtelomeric regions independently in a single channel as a control to quantify barcoding efficiency (Fig. S3, S4 *A*).

We quantified 12 regions that were detected robustly in most cells with a mean of 1.9 ± 0.5 dots (± standard deviation) per cell (Fig. S5, Supporting Text). Consistent with our targeting of 12 distal subtelomeric regions out of a total of 40 distal and proximal subtelomeric regions, we observed that 22.9% of the CRISPR labeled telomere spots corresponded to subtelomeric regions barcoded by DNA seqFISH (Fig. 2 *G*). Similarly, we observed 20.0% of telomere DNA FISH spots corresponded to subtelomere DNA seqFISH spots (Fig. S4 *B*). We note that we do not expect the telomeres and subtelomeres to colocalize perfectly since they can be genomically distant (Table S1, Fig. S4 *A*). We quantified the distribution of the distance between aligned telomeric and subtelomeric spots (Fig. S4 *C*).

From the barcode uniquely assigned to each subtelomeric region, we assigned a unique identity to each tracked region in the live recording. To document the differences of telomeric dynamics from each chromosome, we then analyzed the movements of telomeres assigned to each chromosome (Fig. 2 *H*) and quantified their cumulative square displacements of adjacent time frames as a function of time (Fig. 2 *I*). We also provided multiple quantified traces from additional single cells (Fig. S6).

Based on a calculation of the optical space available in a mammalian nucleus, the single color method could in principle track and identify up to 150 loci (Supporting Text), which could potentially provide a valuable global view of the chromosomes in single cells.

However, there are a few key technological bottlenecks preventing large numbers of loci to be imaged in this fashion. Firstly, targeting non-repetitive regions requires the delivery of a substantial number of distinct sgRNAs to cells. Future work will be focused on ameliorating this limitation. As an alternative to reduce the number of sgRNAs, sets of sgRNAs targeting region-specific repetitive DNAs (10) can be used, while adjacent non-repetitive unique regions or repeat regions themselves can be targeted by DNA seqFISH. Secondly, physical interactions of distinct loci during the live tracking can prevent accurate position tracking, which can be avoided by using multicolor CRISPR labeling (5-10). However, long-term tracking (i.e. beyond a cell-cycle) can be difficult due to the large scale rearrangement and crossovers of chromosomes during mitosis. Lastly, DNA FISH signals can be improved with a robust signal amplification method such as single molecule hybridization chain reaction (smHCR) (12,13) or alternative DNA FISH methods such as CASFISH (14) to increase the detection efficiency.

The key idea in our work is separating the tasks of dynamic tracking of chromosomal loci and the unique identification of these loci. Previous works in multiplexed CRISPR imaging tried to accomplish both goals at the same time, which requires orthogonal Cas9 systems and multiple fluorophores for live imaging. In our approach, we use a single color channel to first track the motion of the chromosomal loci and then use highly multiplexed DNA seqFISH to identify the loci. In addition to the original seqFISH implementation (11), this strategy is another manifestation of the “noncommutative” approach (15) to experimental design that breaks experimental goals into distinct tasks and combines them to accomplish what cannot be easily achieved in a single experimental step. Our method combines advantages of CRISPR labeling and seqFISH for multiplexed live cell detection of genomic loci. Finally, we note that our method can also be combined with sequential RNA FISH (11,12,16) and immunofluorescence to correlate transcriptional and epigenetic states of individual cells with spatiotemporal chromosomal organization in a highly multiplexed manner.

## AUTHOR CONTRIBUTIONS

All authors reviewed and contributed to the writing of the manuscript. L.C., L.S.Q. and Y.T. designed the project. Y.T. and S.H. performed experiments. S.S. wrote analysis codes and Y.T. and S.S. performed data analysis. L.C. supervised the project.

## ACKNOWLEDGMENTS

We thank James Linton for kindly providing cell line and plasmids, and Eric Lubeck for help with probe designing. Y.T. is supported by a graduate fellowship Nakajima Foundation. L.C. is supported by Allen Distinguished Investigator award, and NIH U01-EB 021240-01.

## SUPPORTING CITATIONS

Reference (17,18) appear in the Supporting Material.

## MATERIAL AND METHODS

### Probe design and synthesis

Telomere 59-nucleotide (nt) probe from Integrated DNA Technology (IDT) was designed with a 35-nt targeting sequence at the 3′ end, a 20-nt adapter sequence for binding of a dye-coupled adapter probe, and a 4-nt spacer in between. Subtelomere probes were designed and generated based on array-based oligopool synthesis with enzymatic amplifications (1,2) explained below.

The mm10 mouse genomic sequence (UCSC Genome Bioinformatics) was used to design subtelomere oligonucleotide probe pools in this study. To selectively label subtelomeric genomic regions, 100 kb regions at the end of each chromosome were selected (Table S1). Across those regions, a set of non-overlapping 35-nt probes were designed which suffice several constraints including 40-60% GC content, no more than 5 contiguous identical nucleotides, no “CCCTAA” or “TTAGGG” sequences to exclude the potential binding to telomeres, and at least 2-nt spaces between adjacent probes. Off targets against the mm10 mouse genome were then evaluated using BLAST+. Sequences with 18 or more contiguous bases homologous to other regions in the genome were defined as an off target here, and probes that contained 6 or more of these off targets were initially eliminated. Probes targeting identical subtelomeric regions were then evaluated together, and if the probe sets contained more than 5 off-targets within 1 Mb blocks of the genome, probes were dropped to lower the threshold. If the probe number in one probe set exceeded 400, probes were reduced up to 400 based on GC content. Note that probe sets targeting sex chromosomes were failed to be designed. In addition, proximal telomeres in each chromosome is located adjacent to satellite regions in the mouse genome, so these regions were not used for probe designing. As a result, 19 subtelomere probe sets targeting all mouse autosomes were pooled together in this study (Table S1).

At the 5′ end of the 35-nt probe sets, 20-nt adapter sequences, which are identical in each subtelomere probe set but orthogonal among different probe sets, are attached with a 4-nt spacer in-between. For the array-based oligo library synthesis, universal sequences were attached at either 5′ or 3′ ends. Those sequences included KpnI and EcoRI restriction enzyme sites, 3-nt spacers, and 20-nt forward and reverse primer binding sequences. In total, this subtelomere oligonucleotide probe pool (CustomArray) contained 4709 probes with 117 nucleotides each. Single-stranded DNA probes were generated from this array-based oligonucleotide pool with limited cycle PCR, *in vitro* transcription, reverse transcription, and restriction enzyme digestion of primer binding sites.

### Cell culture and cell line construction

E14 cells (E14Tg2a.4) from Mutant Mouse Regional Resource Centers were maintained on gelatin-coated dishes at 37°C with 5% CO_2_ in Glasgow Minimum Essential Medium (GMEM), 10% FBS (HyClone, Thermo Scientific), 2 mM L-glutamine, 100 units/ml penicillin, 100 μg/ml streptomycin, 1 mM sodium pyruvate, 1000 units/ml Leukemia Inhibitory Factor (LIF, Millipore), 1x Minimum Essential Medium Non-Essential Amino Acids (MEM NEAA, Invitrogen) and 50 μM β-Mercaptoethanol as described previously (3). All constructs used in this study were cloned into PiggyBac vectors. The expression vector for dCas9-EGFP from *Streptococcus pyogenes* was constructed by inserting dCas9-EGFP (pSLQ1658 from Addgene) right after the elongation factor 1 alpha (EF1α) promoter. For the guide RNA expression vector, a mouse U6 promoter and sgRNA targeting telomeres were obtained from pSLQ1651 (Addgene). The vector, which contained EF1α-NLS-HA-NLS-hmKO2 (hmKO2 from Amalgaam), was also constructed and used for cell identification before the live tracking. Transfections were performed with FuGENE HD Transfection Reagent (Promega), and the cells were selected with G418 (Thermo Scientific) and puromycin (Thermo Scientific) sequentially. After the selection, single clones were isolated manually, and stable labeling of telomeres was verified by imaging.

### Live cell imaging

Cells were plated on fibronectin-coated 24-well glass bottom plates (MatTek) for 2 h, prior to the live imaging. The microscope (Nikon Eclipse Ti-E) was equipped with a CCD camera (Andor iKon-M 934), a 60x oil objective lens (Nikon NA 1.40) and a stage-top incubator held at 37°C. Snapshots of dCas9-EGFP were acquired with 10 μm z-stacks stepping every 0.5 μm at 15 time points over 6 min. Note that each time point shown in the figure and movie was the starting time of the z-stacks. The Perfect Focus system of the microscope was used to automatically correct focus drift during imaging. Image acquisition was controlled with Micro-Manager software.

### DNA FISH hybridization and imaging

Immediately after the live cell imaging, cells were fixed in 4% formaldehyde for 10 min at room temperature, washed three times with 1x PBS, and imaged in an anti-bleaching buffer consisting of 20 mM Tris-HCl, 50 mM NaCl, 0.8% glucose, saturated trolox, 0.5 mg/ml glucose oxidase, and catalase at a dilution of 1/1000 (Sigma C3155). Cells were then permeabilized with 70% ethanol at −20 °C overnight. The following day, cells were treated with a prechilled solution of methanol and acetic acid at a 4:1 ratio at room temperature, and then with 0.1 mg/ml RNaseA (Thermo Scientific) for 1 h at 37°C. Samples were then washed and dried with 1x PBS, 70% ethanol and 100% ethanol. The samples were then heated for 10 min at 95°C in 70% formamide and 2x SSC. Cells were hybridized with the telomere and the subtelomere probe pool for 2 days at 37°C, where the final concentration of each probe was estimated as 10 nM in nuclease free water with 50% formamide, 2x SSC and 0.1 g/ml dextran sulfate. After incubation with the probes, cells were washed three times in 50% formamide, 0.1% Triton-X 100 and 2x SSC at room temperature, and hybridized with 20-nt adapter probe sets coupled to Alexa 594, 647 (Lifetech), Cy3B or Cy7 (GE Healthcare) at 10 nM final concentration for at least 1 h at room temperature in nuclease free water with 30% formamide, 2x SSC and 0.1 g/ml dextran sulfate. Cells were washed three times in 30% formamide, 0.1% Triton-X 100 and 2x SSC at room temperature, stained with DAPI and imaged in anti-bleaching buffer.

### Probe displacement and re-hybridization

Following the imaging, cells were washed with 2x SSC, incubated in 70% formamide and 2x SSC for 30 min at room temperature for probe displacement, and then washed three times with 2x SSC. To check the probe displacement, cells were then imaged with all imaging channels in anti-bleaching buffer. Samples were re-hybridized with another set of adapter probes according to the conditions described above, stained with DAPI again and imaged in anti-bleaching buffer.

Four rounds of hybridizations were carried out in this study. The first three rounds of hybridizations were used to barcode 18 subtelomeric regions, and the final round was used to label telomeres and also to verify the identities of 3 subtelomere barcodes by reading out 3 subtelomeric regions with each region assigned to a single imaging channel.

### Data analysis

Data analysis was carried out using ImageJ, MATLAB and Python. Each analysis is detailed below.

### Point tracking

Cells were segmented manually using the ImageJ ROI tool. The background was subtracted from the time-lapse images using ImageJ’s rolling ball background subtraction algorithm with a radius of 3 pixels. This processing was also used for Movie S1. The points for linking in each time point were found in 3D using a LOG filter with subsequent local maxima finding. The threshold for local maxima finding was set using Otsu’s method for the first frame and adjusted slightly for subsequent frames such that the number of dots detected only varied by less than 5%. These points were linked into trajectories using the SimpleTracker function available on the MATLAB file exchange with ‘MaxLinkingDistance’ set to 5 and ‘MaxGapClosing’ set to 0. Any trajectory that did not have a point in all frames was discarded. Every point in every remaining trajectory was then fit with a 2D gaussian function using the autoGuassianSurf function available on the MATLAB file exchange to obtain the subpixel location of the point. Each track was then assigned to a segmented cell. The calculated trajectories were then corrected to remove the motion of the cells and the microscope by subtracting the mean displacement of all points in a cell from each point in the cell for each time point.

For each trajectory, the cumulative square displacement of adjacent frames (CSD) as a function of time was calculated as

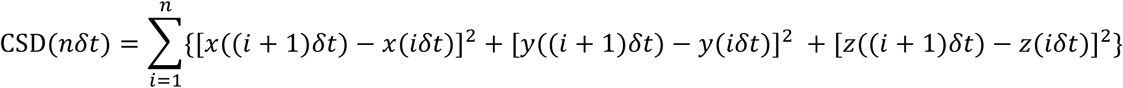

where *n* is the number of frames, *δt* is the time interval between two adjacent frames (25 s), *x*(*t*), *y*(*t*) and *z*(*t*) are the coordinates at time *t*.

### Image processing for barcoding

Basic flow of the image processing for barcoding followed our recent study (2). To remove the effects of chromatic aberration, multispectral beads were first used to create geometric transforms to align all fluorescence channels using MATLAB’s fitgeotrans function. Next, the background illumination profile of every fluorescence channel was mapped using a morphological image opening with a large structuring element on a set of images of an empty coverslip. The median value of every pixel for every channel of opened images was divided by the maximum value to find the division factor of every pixel in every channel. The images were corrected using the resulting intensity map and finally the images were transformed to remove chromatic aberrations. The background signal was then subtracted using the ImageJ rolling ball background subtraction algorithm with a radius of 3 pixels.

### Image registration

The processed images were registered by first taking a maximum intensity projection along the z direction in each channel. All of the maximum projections of the channels in a single hybridization were then collapsed, resulting in 3 composite images containing all the points in a particular round of hybridization. Each of these composite images of hybridizations 2-3 were then registered to hybridization 1 using a normalized cross-correlation algorithm with the position of the maxima of the cross-correlation signifying the translation factor to align hybridizations 2-3 to hybridization 1. MATLAB’s normxcorr2 function was used to accomplish this task. Cross-correlation between the DAPI images was used to register the final control hybridization to the barcoding hybridizations.

### Barcode calling

The potential DNA FISH signals were then found by LOG filtering the registered images and finding points of local maxima above a specified threshold value found by inspection of the accuracy of dots found at a particular threshold value. Once all potential points in all channels in all hybridizations were obtained, dots were matched to potential barcode partners in 3D with all other hybridizations using a √6 pixel search radius (1 or 2 pixel per one direction) to find symmetric nearest neighbors within the given radius. Barcode words were created by seeding the search with points from each hybridization. Point combinations that constructed only a single barcode with a given seed were immediately matched to the on-target barcode set. For points that matched to construct multiple barcodes, first the point sets were filtered by calculating the residual spatial distance of each potential barcode point set and only the point sets giving the minimum residuals were used to match to a barcode. If multiple barcodes were still possible, the point was matched to its closest on-target barcode with a hamming distance of 1. If multiple on-target barcodes were still possible, then the point was dropped from the analysis as an ambiguous barcode. This procedure was repeated using each hybridization as a seed for barcode finding and barcode words that were called uniquely in all hybridizations were used in the analysis. The location of these points then signified the corresponding chromosome locations.

### Dot matching

CRISPR labeled dots at the last frame of the movie and after the fixation, subtelomeric dots by DNA seqFISH and telomeric dots by DNA FISH were matched by using the same matching algorithm described in the barcode calling section, with a small difference of using 6 pixels in xy. In addition, subtelomeric dots by DNA seqFISH and subtelomeric dots by single color DNA FISH readouts in hybridization 4 were matched by using the same algorithm with more stringent matching condition of within 3 pixels. Note that cells detected with more than 10 CRISPR labeled spots at the last frame of the movie were further analyzed due to the heterogeneity of CRISPR labeling efficiency in single cells, and only cells within center fields of view were analyzed to minimize the effect of uneven illumination.

### Number of telomeric and subtelomeric spots

Based on the cell cycle distribution in a mES cell population, we estimated the detection efficiency of telomeric and subtelomeric spots. Typical cell cycle distribution of mES cells is 20% cells in G1, 50% cells in S and 30% cells in G2/M phase (4). Given the number of chromosomal loci is 2 in G1, 3 in S and 4 in M/S2 phase, the number of spots expected per each region is 3.1 per cell. We observed 33.5 ± 13.8 and 40.6 ± 13.8 (mean ± standard deviation) CRISPR labeled dots per cell in live (last frame of the movie) and fixed cells, which can be estimated as 27.0 ± 11.1% and 32.7 ± 11.1% detection efficiency of telomeric spots. This indicates a relatively low efficiency of labeling in our experiment, which can be improved with further cell line engineering as shown in previous publications. Note that we detected more CRISPR labeled spots in fixed cells compared to those in live cells because of longer imaging exposure time for fixed cells. We also note that we used exposure times that allowed us to track CRISPR labeled loci over time without significant photobleaching. However, we still observed that the number of spots detected above the threshold decreased during the time-lapse movie, because of photobleaching (Fig. S1 *A*). Similarly, DNA FISH of the telomeres showed 32.5 ± 7.6 dots per nuclei and 26.2 ± 6.1% detection efficiency of telomeric spots. The relatively low colocalization efficiency (49.1%) of telomeric spots by CRISPR labeling and DNA FISH (Fig. S4 *B*) can be caused by the low labeling efficiencies estimated above.

On the other hand, from our barcoding results, the average number of subtelomeric spots per cell was 1.9 ± 0.5, and the DNA seqFISH efficiency of subtelomeric regions can be estimated as 61.3 ± 16.1%.

### Optical space estimation in nucleus

Optical space for single-color CRISPR labeling in a single nucleus can be estimated based on our recent study (2). The estimation is calculated as

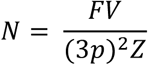

where *N* is the maximum number of unambiguous CRISPR labeled spots in a single nucleus, *F* is the number of channel used for CRISPR imaging, *V* is the volume of a single nucleus in microns, *p* μm is the physical size of a pixel and *Z* μm is the resolution in the z direction. In our experimental condition, a single nucleus can accommodate at least 1000 CRISPR labeled spots by applying a single fluorescent channel, the physical pixel size 0.3 μm, z resolution 0.5 μm and the volume of mES cell nucleus as 10 μm × 10 μm × 5 μm.

The number of CRISPR labeled spots, which can be uniquely identified by DNA seqFISH in a single nucleus, are reduced with the optical space constraint arising from the incomplete colocalization between two labeling methods. Under such conditions, the estimation is updated as

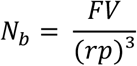

where *N_b_* is the maximum number of unambiguous CRISPR labeled spots identified by DNA seqFISH in a single nucleus and *r* is the maximum searching pixel size per single direction for dot matching. Given the same assumption above with 5 pixel diameter search, a single nucleus can accommodate around 150 CRISPR labeled spots that can be uniquely identified by DNA seqFISH. Note that the number of uniquely identified loci can be linearly scaled up with the increase of fluorescent channels available for the CRISPR imaging.

**Table S1:** Subtelomeric region coordinates in mm10 mouse genome, number of primary probes, sequence gap between telomere and targeted subtelomeric region, and barcoding color combinations used in this study. Sequence gap was calculated as the length between distal telomere coordinate annotated and the most adjacent subtelomeric probe in each chromosome. Due to the off targets, chromosome 2 probe set was not included in the DNA seqFISH. Cy3B, Alexa 594, 647 and Cy7 dye coupled adapter probes correspond to the numbers 1, 2, 3 and 4 in the last 4 columns. Finally, 12 subtelomeric regions (chr1, 3, 5, 6, 7, 9, 13, 15, 16, 17, 18 and 19) were read out robustly.

| Chrom | Start | End | Strand | Probe number | Sequence gap (bp) | hyb1 | hyb2 | hyb3 | hyb4 |
| --- | --- | --- | --- | --- | --- | --- | --- | --- | --- |
| chr1 | 195271955 | 195371955 | + | 189 | 8371 | 3 | 3 | 3 |  |
| chr2 | 181913208 | 182013208 | + | 159 | 11144 |  |  |  |  |
| chr3 | 159839664 | 159939664 | + | 400 | 638 | 2 | 3 | 2 |  |
| chr4 | 156208115 | 156308115 | + | 400 | 152242 | 4 | 2 | 2 |  |
| chr5 | 151634668 | 151734668 | + | 400 | 268 | 1 | 1 | 3 |  |
| chr6 | 149536530 | 149636530 | + | 272 | 50346 | 1 | 3 | 1 |  |
| chr7 | 145241443 | 145341443 | + | 400 | 542 | 1 | 3 | 2 |  |
| chr8 | 129101212 | 129201212 | + | 100 | 115103 | 4 | 1 | 3 |  |
| chr9 | 124395094 | 124495094 | + | 341 | 6138 | 1 | 1 | 2 | 3 |
| chr10 | 130494977 | 130594977 | + | 105 | 752 | 3 | 4 | 2 |  |
| chr11 | 121832542 | 121932542 | + | 186 | 50093 | 2 | 1 | 2 |  |
| chr12 | 119929006 | 120029006 | + | 400 | 688 | 4 | 1 | 2 | 1 |
| chr13 | 120221623 | 120321623 | + | 126 | 960 | 3 | 1 | 2 |  |
| chr14 | 124702228 | 124802228 | + | 179 | 727 | 4 | 2 | 1 |  |
| chr15 | 103843669 | 103943669 | + | 187 | 112 | 2 | 1 | 1 |  |
| chr16 | 98007752 | 98107752 | + | 115 | 8793 | 1 | 2 | 2 |  |
| chr17 | 94787255 | 94887255 | + | 105 | 726 | 2 | 1 | 3 |  |
| chr18 | 90502623 | 90602623 | + | 400 | 1355 | 1 | 2 | 1 | 2 |
| chr19 | 61231550 | 61331550 | + | 245 | 611 | 2 | 2 | 2 |  |

**Figure S1:**
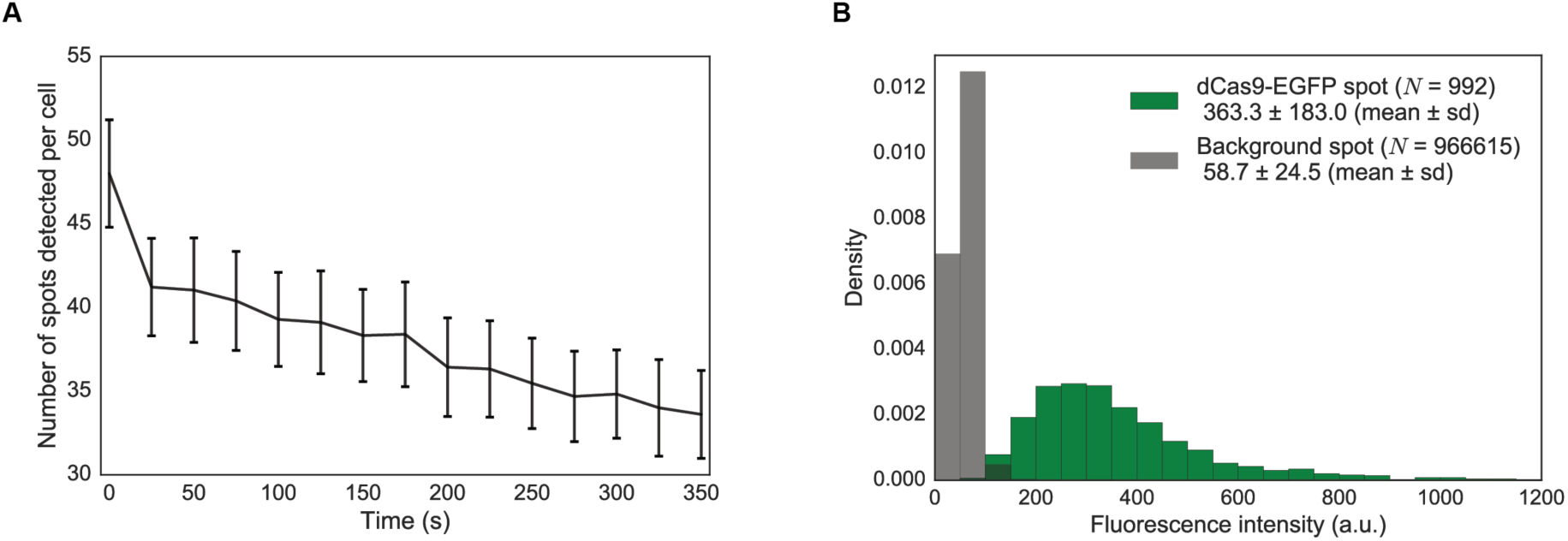
Number of telomeric spots detected per cell during the movie and their fluorescence intensity. (*A*) Decrease of number of telomeric spots detected per cell during tracking due to photobleaching. The threshold used for ‘CRISPR live cells’ in Figure 2F was used in all time points. The data are displayed as mean ± sem with 28 cells. (*B*) Distribution of fluorescence intensity of detected dCas9-EGFP spots and background spots at the last frame of the movie. The intensity of dCas9-EGFP spots were detected as a maximum intensity within 3×3 pixels, and the intensity of background spots were collected after eliminating those 3×3 pixels.

**Figure S2:**
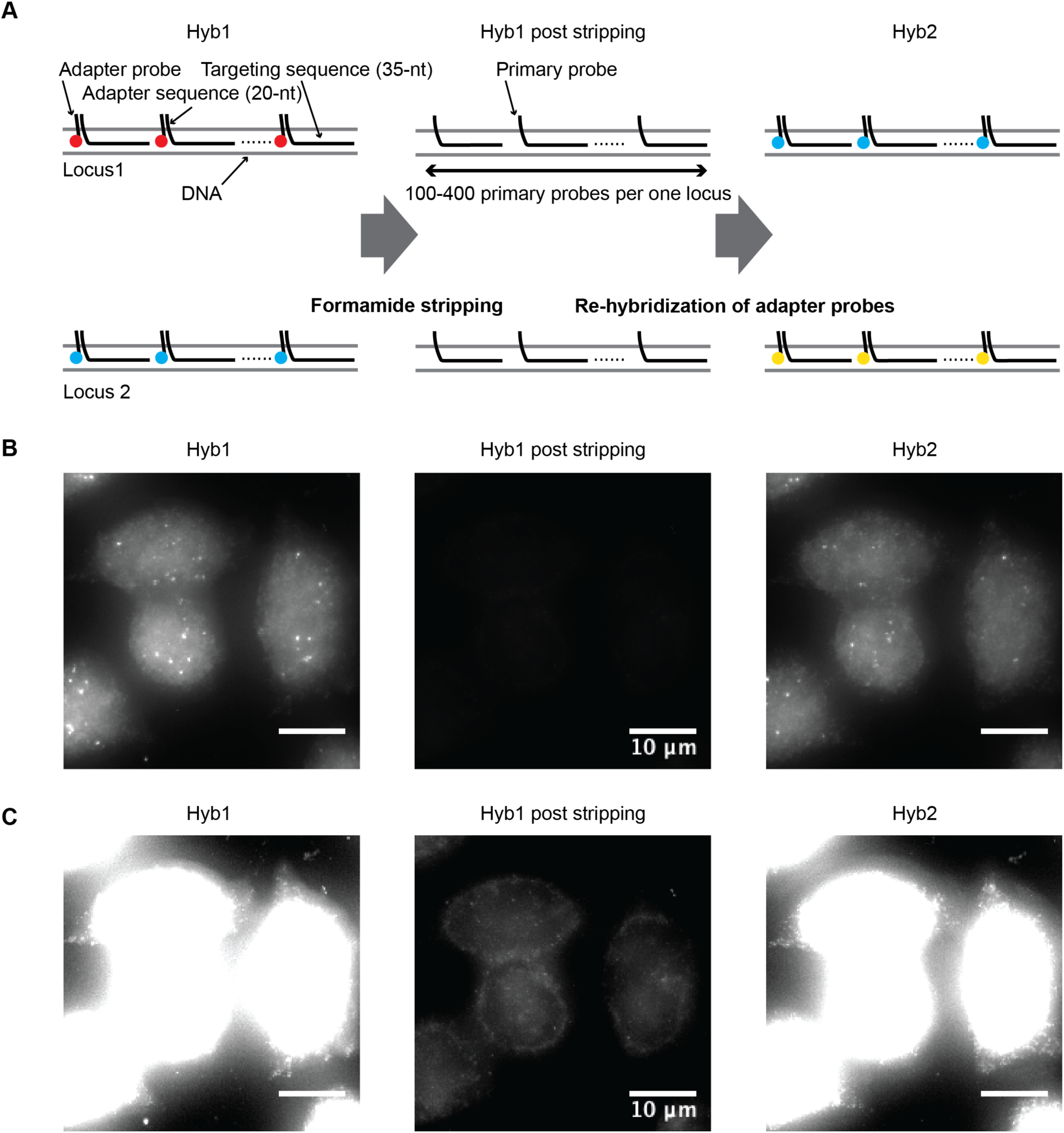
Probe displacement and re-hybridization. (*A*) Schematic of probe displacement and re-hybridization with two loci. (*B, C*) From left to right: first round of adapter probe set hybridization, stripped cells after probe displacement with the formamide stripping method, and second round of hybridization containing different adapter probe combinations from the first hybridization in mES cells. All images are maximum intensity projections of a z-stack with Cy3B adapter probe sets, and displayed at two contrast levels (*B* and *C*) to show the completeness of stripping.

**Figure S3:**
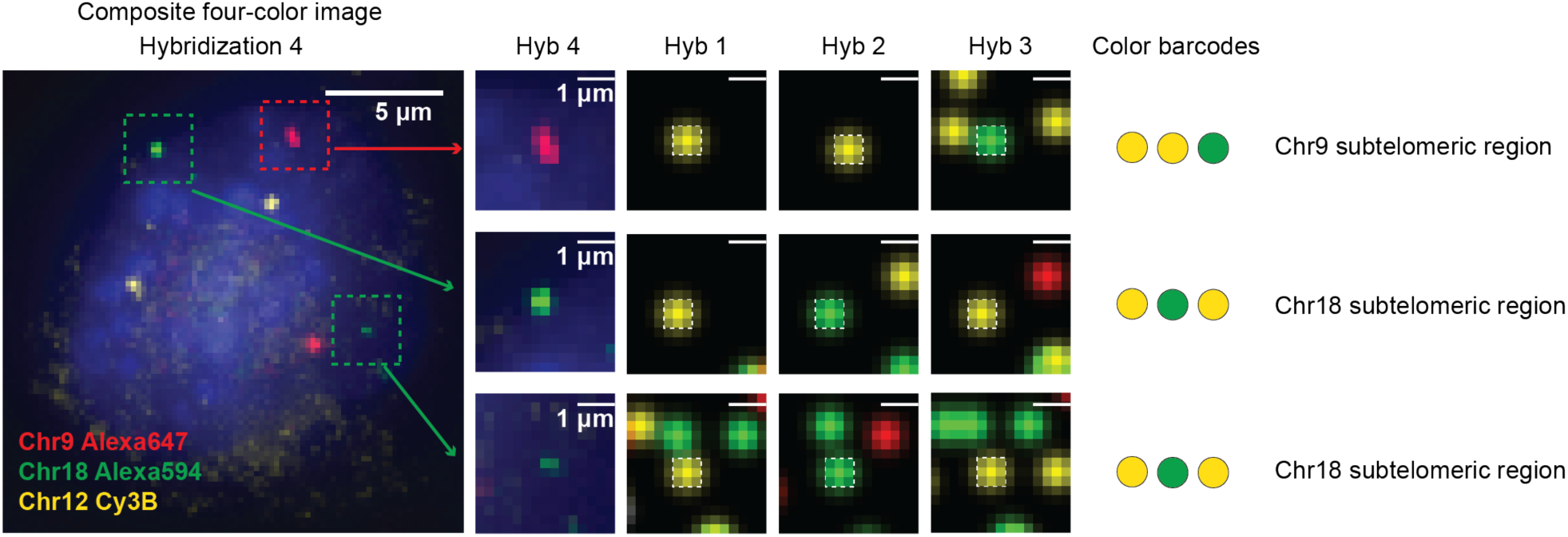
Comparing single color DNA FISH readouts (hybridization 4) and DNA seqFISH color barcoding (hybridizations 1-3) in mES cells. Images are maximum projections of a z-stack. Boxed regions in the left figure are magnified and corresponding regions in hybridizations 1-4 are displayed. Each color represents Alexa 647 (red), Alexa 594 (green), Cy3B (yellow) and DAPI (blue), respectively. Images with hybridizations 1-3 are digitized based on the barcode calling results. Dots appearing in hybridizations 1-3 images other than the dots colocalized to the hybridization 4 are dots corresponding to other barcodes or nonspecific binding. We observed that with the chromosome 9 subtelomeric region, 78.7% of the single color labeled loci in the fourth hybridization (53 spots analyzed) colocalized with the barcoded loci (53 spots analyzed), whereas with the chromosome 18 subtelomeric region, 73.7% of the single color labeled loci in the fourth hybridization (92 spots analyzed) colocalized with the barcoded loci (75 spots), indicating barcodes decoded efficiently in our experiments. Note that the chromosome 12 subtelomeric region was excluded from this analysis due to the insufficient signal from the Cy7 dye in DNA seqFISH.

**Figure S4:**
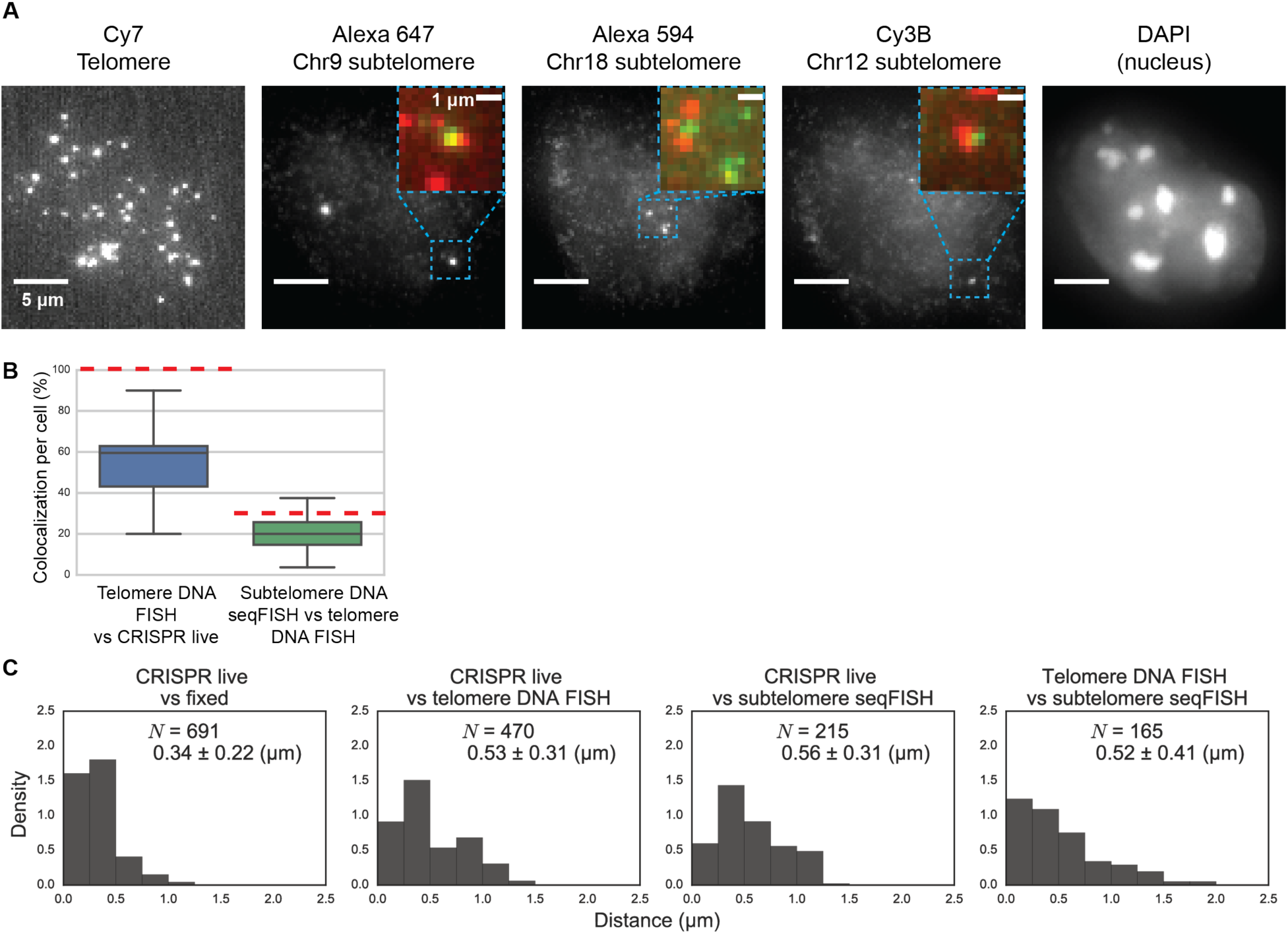
Colocalization between telomeric and subtelomeric spots and their distribution in mES cells. (*A*) Images are maximum intensity projections of a z-stack of fluorescence images corresponding to the fourth hybridization of the DNA seqFISH. The boxed regions are magnified, and telomeric (red) and subtelomeric (green) regions are merged. Note that telomeric and subtelomeric regions do not colocalize perfectly because targeted telomeric regions are non-unique repetitive regions whereas targeted subtelomeric regions are adjacent unique regions over a range of 100 kb. Note that sequence spaces between telomeric and subtelomeric regions are provided in Table S1. (*B*) Comparing colocalization percentage of spots detected per cell. Red dashed lines represent expected colocalization percentage per cell. (*C*) Distribution of xy-distance between aligned telomere CRISPR spots, subtelomere DNA seqFISH spots and telomere DNA FISH spots. Mean and standard deviation of the distance under each condition were provided.

**Figure S5:**
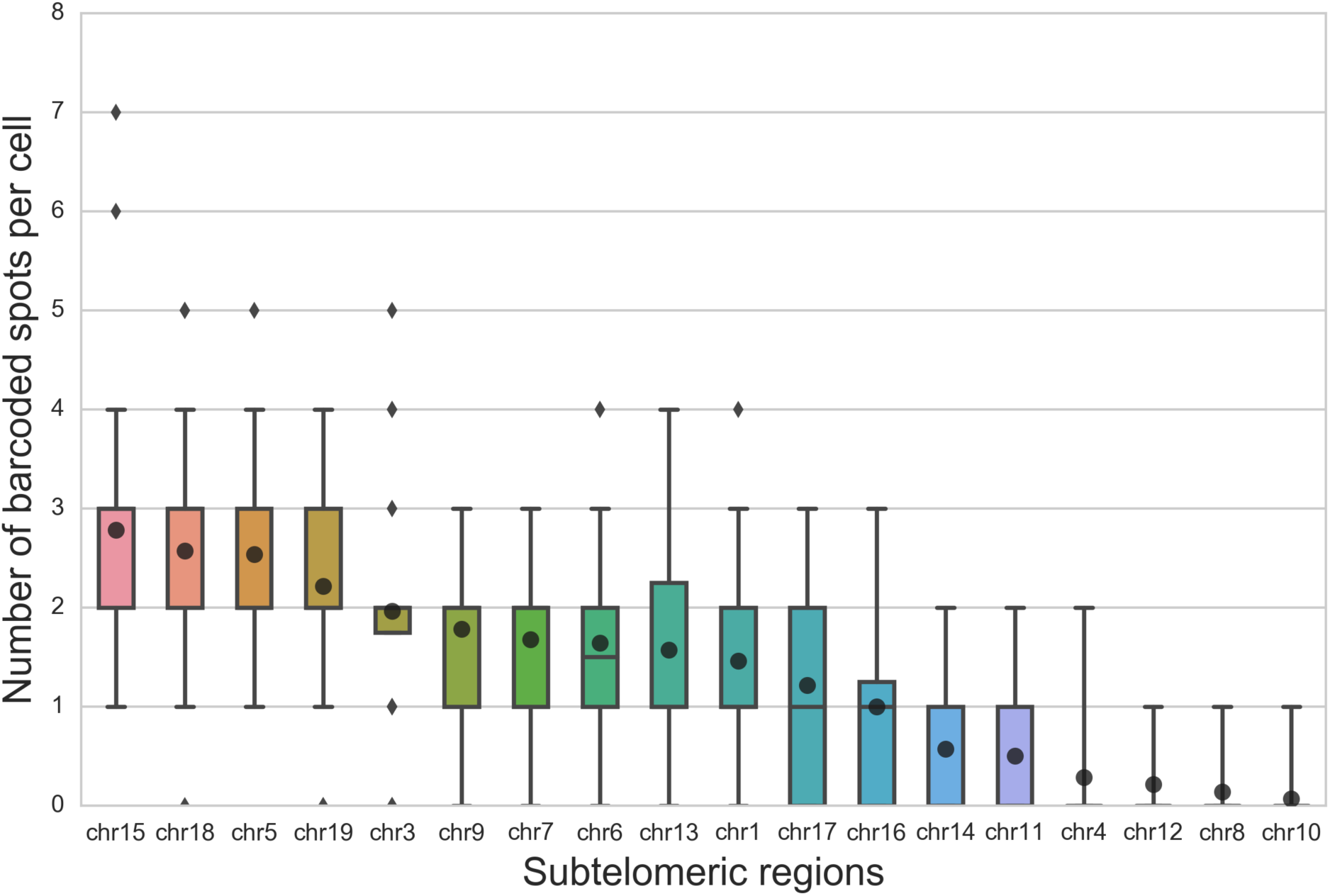
Number of subtelomeric spots per cell resolved by the color barcoding with three rounds of hybridizations. In total, 678 subtelomeric spots in 28 cells were analyzed. Black circles represent mean number of spots per cell. Due to the low detection efficiencies, 6 subtelomeric regions (chr14, chr11, chr4, chr12, chr8 and chr10) were excluded from the analysis. This could be caused by inefficient binding of primary probe sets or insufficient signal from Cy7 fluorophores as 5 out of those 6 subtelomeric regions contained Cy7 in their code. On average, the number of subtelomeric spots per cell was 1.9 ± 0.5 (mean ± standard deviation).

**Figure S6:**
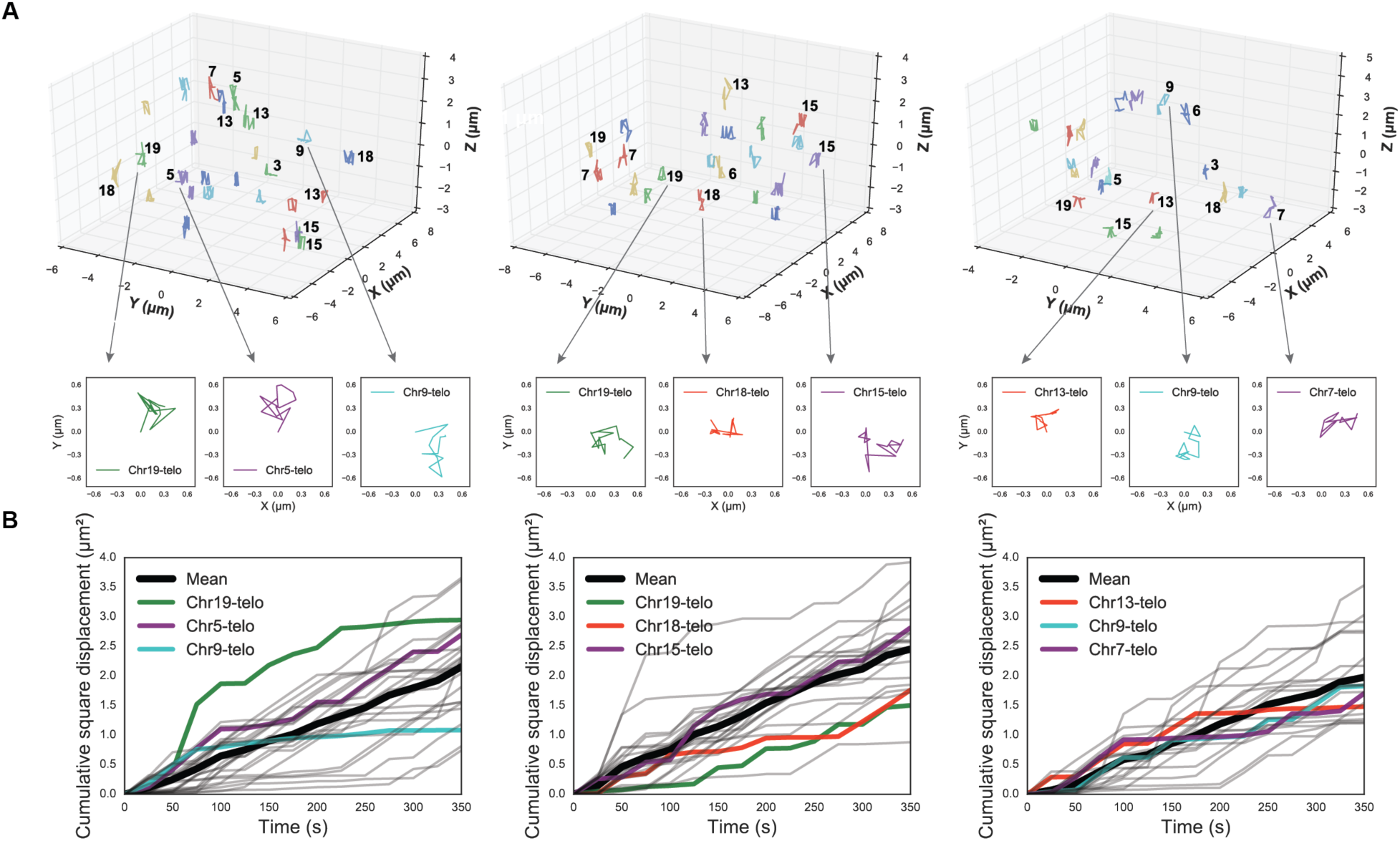
Quantified trajectories of telomeric loci from three additional single cells. (*A*) In those cells, 26, 23 and 20 trajectories were detected from CRISPR imaging, and 13, 9 and 9 of these trajectories (from left to right) were uniquely assigned to particular chromosomes based on the subtelomere color barcodes. Trajectories of three loci per cell were also highlighted as xy projections (inset). Projected trajectories start from (0.0, 0.0). (B) Cumulative square displacement traces as a function of time. Those traces were obtained from the three single cells shown above. Three projected loci per cell (*A* inset) were shown as colored traces.

## Supporting Movies

**Movie S1**: Live imaging of telomeres in mES cells using the CRISPR labeling. Cells shown in Fig. 2*B* are presented. Images are maximum intensity projections of a z-stack of fluorescence images in each frame. Note that cell and stage movements are not calibrated in this movie. Scale bar represents 10 μm.

